# *Enterococcus faecium sagA* mutants have cell envelope defects influencing antibiotic resistance and bacteriophage susceptibility

**DOI:** 10.1101/2025.07.21.665895

**Authors:** Garima Arya, Pavan Kumar Chodisetti, Juliel Espinosa, Brian C. Russo, Howard C. Hang, Breck A. Duerkop

## Abstract

*Enterococcus faecium* is a Gram-positive bacterium that is resident to the intestines of animals including humans. *E. faecium* is also an opportunistic pathogen that causes multidrug resistant (MDR) infections. Bacteriophages (phages) have been proposed as therapeutics for the treatment of MDR infections, however, an obstacle for phage therapy is the emergence of phage resistance. Despite this, the development of phage resistance can impact bacterial fitness, thus, understanding the molecular basis of fitness costs associated with phage resistance can likely be leveraged as an antimicrobial strategy. We discovered that phage resistant *E. faecium* harbor mutations in the cell wall hydrolase gene *sagA*. SagA cleaves crosslinked peptidoglycan (PG) involved in PG remodeling. We show that mutations in *sagA* compromise *E. faecium* PG hydrolysis rendering them sensitive to β-lactam antibiotics. *sagA* mutants have cell envelope integrity defects, increased cellular permeability, and aberrant distribution of penicillin binding proteins. This corresponds to a growth defect where cells have abnormal division septa, membrane blebbing, and the formation of mini cells. The dysregulation of the cell envelope in *sagA* mutants alters the binding of phages to the *E. faecium* cell surface. Our data support a model where phage infection of *E. faecium* requires phages to localize to sites of peptidoglycan remodeling at the cell poles and division septa. Our findings show that by altering the function of a single PG hydrolase, *E. faecium* loses intrinsic β-lactam resistance. This indicates that phage therapy could help revive certain antibiotics when used in combination.

## Introduction

*Enterococci* are intestinal Gram-positive bacteria that cause severe systemic infection (1–4). Enterococcal bloodstream infections are associated with high mortality, and this has increased following the COVID-19 pandemic (5, 6). Among medically relevant enterococci, *Enterococcus faecium* and *Enterococcus faecalis* are of particular concern because they are hospital-adapted and cause nosocomial infections. Nosocomial *E. faecium* infections are problematic due to the organism’s intrinsic and acquired resistance to diverse antibiotics, including β-lactams, aminoglycosides, and vancomycin (7–9). Notably, prior exposure to cephalosporins, a type of β-lactam, is a leading risk-factor for *E. faecium* infection (10, 11). Additionally, a hallmark feature of *E. faecium* is evolved antibiotic resistance through the acquisition of foreign DNA via horizontal gene transfer, which has directly contributed to their ability to thrive as a hospital adapted opportunistic pathogen (12, 13).

There is growing interest in developing approaches to limit enterococci from hospital settings, including targeted screening of at-risk patients, elevated hygiene practices, and limiting hospital stays (14–16). Aside from interventions to patient care, innovations in therapeutic approaches including the use of bacteriophages (phages – viruses that infect bacteria) for the treatment of multi-drug resistant (MDR) enterococcal infections are gaining traction (17–21). Despite the excitement surrounding the potential of phage therapy, phage predation can lead to phage-resistant bacteria during treatment (22–24). Although phage resistance has the potential to threaten the efficacy of phage therapies, there is mounting evidence that fitness constraints arise in phage resistant bacteria leading to trade-offs such as heightened antibiotic sensitivity, dampening of virulence, and colonization defects (25–29). Considering fitness trade-offs stemming from phage resistance are well-documented and could be exploited therapeutically, we know very little about molecular mechanisms that support these trade-offs.

Our group discovered that *E. faecium* phage resistant isolates are more susceptible to β-lactam antibiotics including ampicillin and ceftriaxone (26). Sensitivity to cephalosporins is the result of mutations in SagA, a hydrolase that cleaves peptidoglycan during cell division (26, 30–33). Here we show that *E. faecium sagA* mutants have compromised cell wall integrity resulting in increased cellular permeability and drug sensitivity. *E. faecium sagA* mutants exhibit diffuse penicillin binding protein (PBP) localization throughout the periphery of the cell. Additionally, *sagA* mutants have cell-wall morphology defects characterized by aberrant division septa, cell envelope protrusions, and mini cell formation that support phage resistance. Phages bind to discrete sites on the cell surface of *E. faecium* that are associated with active peptidoglycan synthesis, whereas, phages are sequestered to cell envelope protrusions and mini-cells in the *sagA* mutant resulting in non-productive phage infection. These findings provide a mechanistic basis for how SagA deficiency leads to β-lactam sensitivity and cell wall defects that support phage resistance. Intrinsic resistance to β-lactams is a common feature of *E. faecium,* and our findings support the idea of revisiting β-lactams as a potential treatment in combination with phage therapy.

## Results

### The peptidoglycan hydrolase domain of *E. faecium* SagA is required for peptidoglycan cleavage and β-lactam resistance

Prior work from our group revealed that *E. faecium* Com12 develops resistance to the phage 9181 by acquiring mutations in the *sagA* gene (26). *sagA* encodes a NlpC/P60 peptidoglycan hydrolase that cleaves crosslinked peptidoglycan (PG) during cell wall remodeling (31, 33). Initially thought to be an essential gene (34), we recovered a variety of point mutations near critical amino acid residues in the NlpC/P60 hydrolytic domain, suggesting that these mutations rendered SagA nonfunctional. To test if these phage mediated mutations in SagA hamper its PG hydrolytic activity, we assessed the in vitro enzymatic activity of *E. faecium* Com12 SagA from two independent phage resistant mutant strains. Enzymatic activity was measured using mutanolysin-digested peptidoglycan from *E. faecium* Com12 and the production of GlcNAc-MurNAc-dipeptide (GMDP) (Fig 1A). We initially chose to focus on two *E. faecium* mutant strains 81R6 and 81R8, which have an F insertion between Y451 and L452 and a G435V substitution in the catalytic domain, respectively. Our attempts to purify the 81R6 SagA mutant version of SagA (F insertion between Y451 and L452) were unsuccessful. Therefore, to test if the position of this amino acid insertion is crucial for the catalytic activity of SagA, we made a version of SagA that has the smaller hydrophobic amino acid leucine inserted at this position. Both SagA mutants have a reduced ability to hydrolyze PG suggesting that point mutations localized to the NlpC/P60 catalytic domain disrupt enzymatic activity (Fig 1B, 1C). Consistent with this observation, the Hang lab recently showed that a chromosomal deletion of *sagA* could be made in the related *E. faecium* strain Com15, further confirming *sagA* to be a non-essential gene (33).

**Fig 1.**
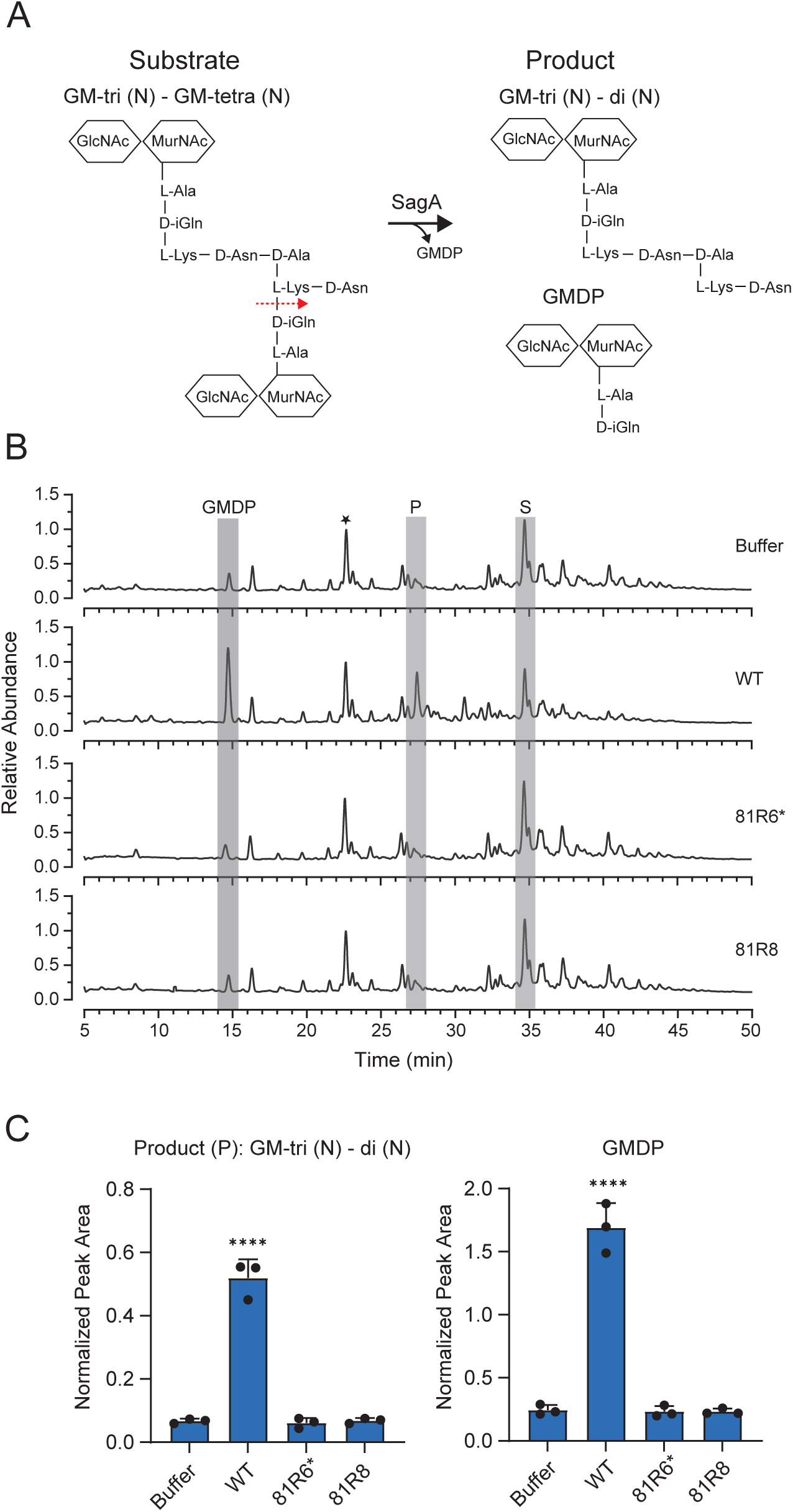
Phage resistant sagA mutants have compromised D, L-endopeptidase activity. **(A)** Schematic showing SagA endopeptidase activity. Substrate denotes the cross-linked muropeptide released following mutanolysin digestion of intact *E. faecium* Com12 PG. SagA mediated cleavage of substrate releases GlcNAc-MurNAc-dipeptide (GMDP) and Product: GM-tri (N) - di (N). **(B)** LC-MS analysis of *E. faecium* Com12 PG incubated with purified SagA, SagAY451/L452-L ins (81R6*), SagA G435V (81R8). Total ion chromatogram was normalized to the intensity of unchanging GM-tri (D-Asn) peak marked with a star in the buffer control. P, Product; S, Substrate **(C)** Quantification of Product and GMDP formation by SagA, SagAY451/L452-L ins (81R6*), SagA G435V (81R8). Extracted ion chromatograms of SagA-treated samples were compared to the buffer control by one-way ANOVA with Dunnett’s post-test (n = 3). ****p < 0.0001. Comparisons with no asterisk had P > 0.05 and were not considered significant.

Our previous work also indicated that *E. faecium* SagA mutants were more susceptible to cell wall-targeting antibiotics and specifically cephalosporins for which *E. faecium* is intrinsically resistant (35). Thus, to test the impact SagA functionality for *E. faecium* resistance to cell wall-targeting antibiotics, we determined minimal inhibitory concentrations (MICs) using microbroth dilution assays against representative β-lactams including 2^nd^(cefuroxime)-, 3^rd^(ceftriaxone)-, 4^th^(cefepime)-, and 5^th^(ceftaroline)-generation cephalosporins, ampicillin, and meropenem, as well as, the cyclic peptide bacitracin which inhibits lipid carriers during PG synthesis (36). The *sagA* mutant strain 81R6 exhibited reduced MICs of at least two-fold or greater for all antibiotics tested (Table 1), with sensitivity the most enhanced for ceftriaxone (32-fold) and cefepime (128-fold). Importantly, ceftriaxone resistance was restored in *E. faecium* 81R6 by complementing *sagA* on a plasmid (Table S1). Together, these observations are consistent with our earlier findings where loss of functional SagA results in antibiotic sensitivity (26, 33) and also extend this phenotype to a diversity of cell-wall targeting antibiotics.

**Table 1.** *sagA* mutant 81R6 have enhanced susceptibility to antimicrobials.

| **Antimicrobial** | ***E. faecium* Com12 MIC (µg/ml)<sup>a</sup>** |  |
| --- | --- | --- |
|  | **Wild type** | **81R6** |
| Cefuroxime | 512 | 128 |
| Ceftriaxone | 32 | 1 |
| Cefepime | 256 | 2 |
| Ceftaroline | 32 | 2 |
| Ampicillin | 4 | 1 |
| Meropenem | 64 | 32 |
| Bacitracin | 256 | 128 |

<sup>a</sup>Median MICs determined from at least three independent replicates

### Loss of SagA function compromises *E. faecium* cell wall integrity

Since SagA is involved in PG hydrolysis that supports symmetric cell division (33), we hypothesized that loss of *sagA* activity would result in compromised cell wall integrity. To test this, we exposed *E. faecium* Com12 and the *sagA* mutant strains 81R6 and 81R8 to 2% sodium dodecyl sulfate (SDS), an anionic detergent that disrupts membranes leading to bacterial cell lysis. When exposed to SDS, wild type *E. faecium* experiences minimal cell lysis measured by the release of cytoplasmic protein (Fig 2A). As a control we first treated *E. faecium* Com12 with lysozyme to weaken the cell wall and then exposed the cells to SDS, which was sufficient to lyse the cells (Fig. 2A). *E. faecium sagA* mutants, when treated with SDS alone, released high amounts of intracellular protein (Fig 2A). Protein release by these mutants in the presence of only SDS was comparable to that of wild type cells treated with lysozyme and SDS. SDS resistance was restored by complementing *sagA* in the mutant strains (Fig 2B). Together, these data indicate that loss of SagA function compromises *E. faecium* cell wall integrity.

**Fig 2.**
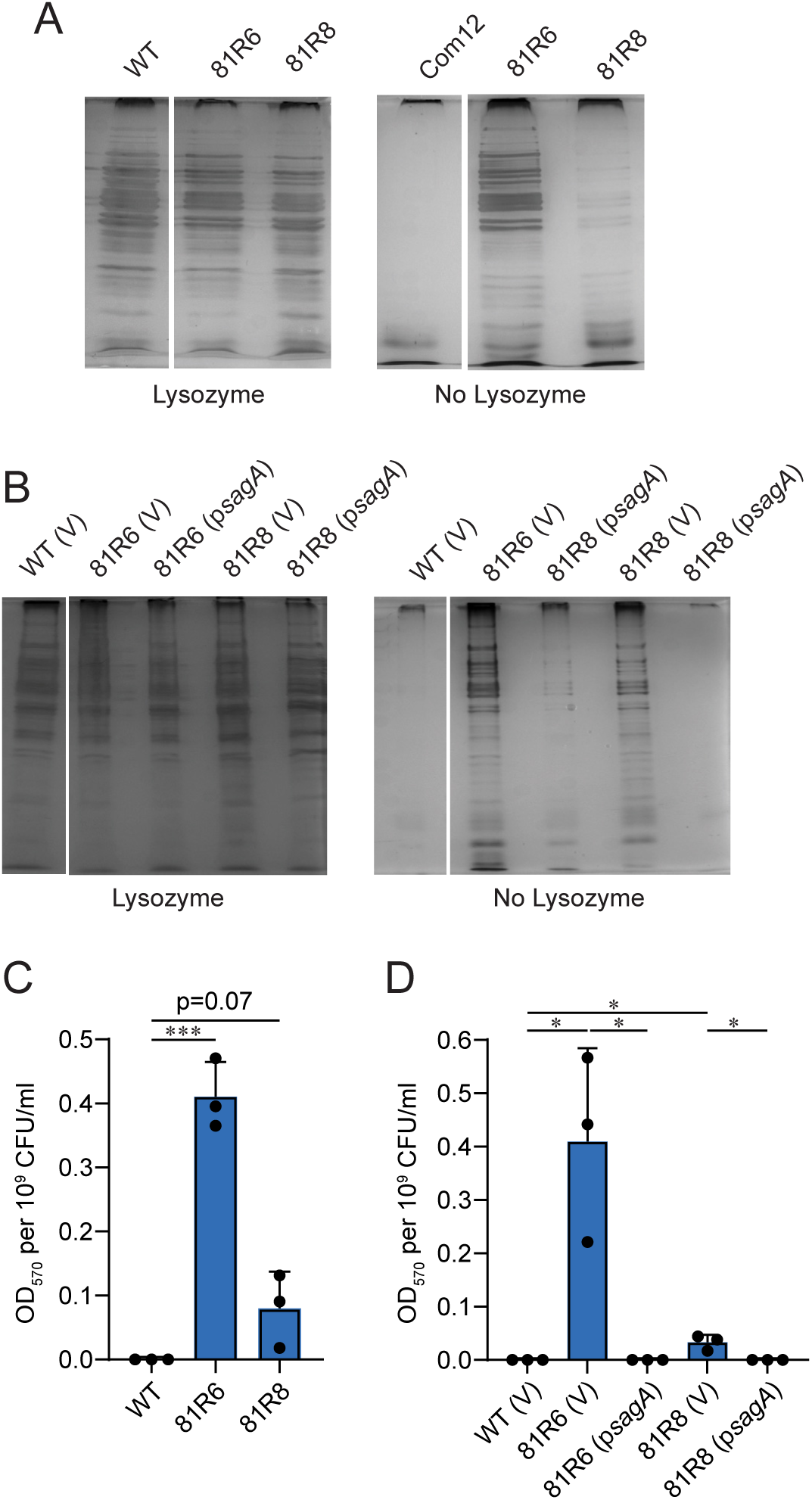
*E. faecium sagA* mutants exhibit compromised cell envelope integrity. **(A-B)** Cell envelope integrity of *E. faecium* Com12 (WT) and *sagA* mutant strains 81R6 and 81R8 strains. Strains carrying plasmid empty pAM401 (V) or pAM401 with Com12 *sagA* (p*sagA*). Exponentially grown cells in BHI were harvested and treated with (+) or without (−) lysozyme, prior to the addition of SDS Laemmli sample buffer. Samples were subjected to SDS-PAGE. Protein bands were visualized by silver staining. **(C-D)** Cell envelope damage for *E. faecium* strains described in panels A and B was assessed by measuring the hydrolysis of CPRG. CPRG hydrolysis was quantified after removing cells and measuring absorbance at 570 nm of the cell supernatant normalized to viable CFU. Data represents average (±SD) from three independent experiments. ND, not detected. p values were calculated using unpaired two-tailed Student’s t test (***p < 0.001; **p < 0.01; *p < 0.03).

Next, we sought to determine if the compromised cell wall integrity in these phage resistant *sagA* mutants leads to altered cell envelope permeability. To test this, we performed a small molecule permeability assay that relies on the accessibility of the substrate chlorophenol red β-D galactopyranoside (CPRG), an otherwise impermeant small molecule, to intracellular β-galactosidase (37). CPRG enters permeable cells and β-galactosidase hydrolyses CPRG into the colored compound chlorophenol red that can be quantified by measuring absorbance at 570 nm. *E. faecium* Com12 with an intact cell envelope is impermeable to CPRG and no hydrolysis occurs (Fig 2C). However, the *sagA* mutants 81R6 and 81R8 hydrolyzed CPRG to varying degrees indicating a defective cell envelope and increased permeability (Fig 2C). Cell envelope impermeability was restored comparable to wild type *E. faecium* when 81R6 and 81R8 were complemented with *sagA* (Fig 2D). These results show that *sagA* mutants have impaired cell wall integrity and increased permeability to small molecules.

### *E. faecium sagA* mutants exhibit increased penetration of β-lactams and disordered penicillin binding proteins

To determine if β-lactams could penetrate the *E. faecium* cell envelope more readily in the *sagA* mutants, we performed a β-lactam penetration assay using Bocillin, a fluorescent BODIPY-labeled version of penicillin, which acylates PBPs similar to ceftriaxone (38). We focused our proceeding work on the *sagA* mutant strain 81R6, as this strain had a significant reduction in the ceftriaxone MIC (26) (Table 1) and experienced maximal damage to the cell envelope (Fig 2C). Compared to *E. faecium* Com12, 81R6 showed increased uptake of Bocillin regardless of the concentration tested (Fig 3A). Because ceftriaxone resistance in *E. faecium* is governed by the low affinity PBPs PbpA(2b) and Pbp5 and the class A PBPs PonA and PbpF (39, 40), increased cell envelope permeability in *E. faecium* 81R6 could lead to rapid PBP acylation by β-lactams elevating ceftriaxone sensitivity. To assess this, we first measured ceftriaxone-mediated acylation patterns of PBPs when competing with Bocilllin. Exponentially grown cells were treated with increasing concentrations of ceftriaxone followed by excess Bocillin. Any PBPs acylated by ceftriaxone will not be labeled by Bocillin, whereas, ceftriaxone non-reactive PBPs will be acylated by excess Bocillin and fluoresce. SDS-PAGE and fluorescent protein band imaging shows that in the absence of ceftriaxone PBPs are readily acylated by Bocillin (Fig 3B). In the presence of ceftriaxone, we did not observe any significant difference in the intensity of five of the Bocillin fluorescent bands (BFB) - BFBs 2-6, between *E. faecium* Com12 and 81R6. However, we observed a slightly less intense band (BFB 1) at the highest concentration of ceftriaxone (64µg/ml) for 81R6 (Fig. 3B). Together, this indicates that the heightened cell envelope permeability of 81R6 results in limited to no substantial impact on PBP acylation patterns.

**Fig 3.**
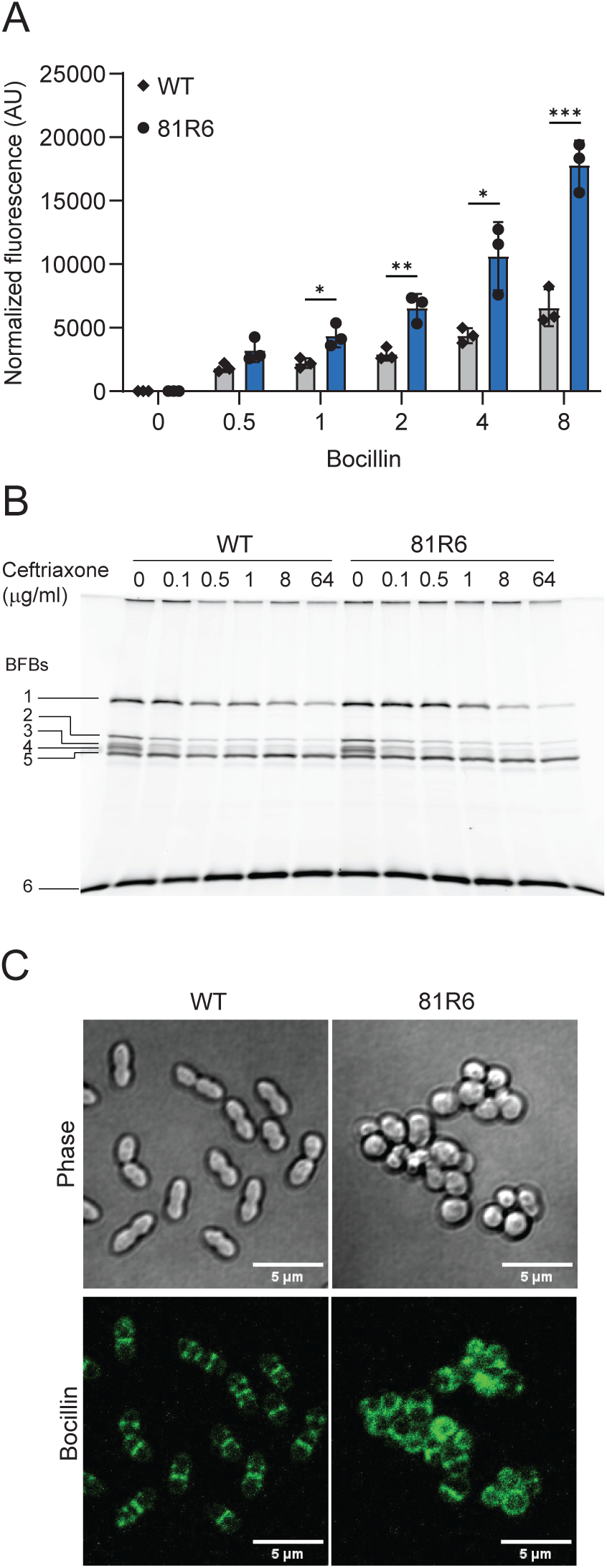
In vivo acylation of PBPs in *E. faecium sagA* mutants. **(A)** The *E. faecium sagA* mutant 81R6 is more reactive to Bocillin uptake. Fluorescence was measured at 488 nm and normalized to optical density (OD_600_) of the samples. Data represents average (±SD) from three independent experiments. ND, not detected. p values were calculated using multiple comparisons t test with adjusted p values using Holm-Šidák method. ***p < 0.002; **p < 0.007; *p < 0.02 **(B)** Bocillin fluorescently labeled bands (BFBs) in the presence of ceftriaxone in membrane fraction of *E. faecium* strains. Membrane fraction without ceftriaxone treatment was taken as control. BFBs (labeled as #1 to 6) were separated within SDS-PAGE gels and bands were detected by imaging at 488 nm. Experiment was repeated at least three times. **(C)** Bocillin staining of *E. faecium* to show PBP distribution in the *sagA* mutant 81R6 compared to Com12 (WT) cells.

For proper cell wall synthesis, PG hydrolases work in close association with PBPs (41, 42). SagA is a key PG hydrolase involved in PG remodeling, so it is possible that *sagA* mutations could alter the distribution of PBPs within the cell. To test this, we stained actively dividing *E. faecium* Com12 and 81R6 with Bocillin and visualized PBP localization using fluorescence microscopy. Wild type cells show a diplococcus morphology with Bocillin labelling at the equator and newly formed division septa (Fig 3C). In contrast, 81R6 cells showed mis-localized Bocillin staining throughout the periphery of the cell along with random sites of dense Bocillin accumulation (Fig. 3C). These data indicate that PBPs are disorganized in the absence of functional SagA and fail to localize properly during cell division.

### Peptidoglycan remodeling is altered in *E. faecium sagA* mutants

SagA supports cell envelope integrity in *E. faecium*, thus we hypothesized that mutating *sagA* would impair PG synthesis. To assess this, we analyzed *E. faecium* Com12 and 81R6 for their ability to incorporate 3-[[(7-Hydroxy-2-oxo-2H-1-benzopyran-3-yl)carbonyl]amino]-D-alanine (HADA), a fluorescent analog of D-alanine. Using fluorescence microscopy, we observed that *E. faecium* Com12 cells showed tri-septal HADA staining at the equator and septa, whereas, 81R6 showed uniform distribution of HADA around the periphery of the cell, suggesting a defect in PG synthesis (Fig 4A). Complementation of 81R6 with *sagA* restored the tri-septal HADA staining pattern (Fig 4A). Transmission electron microscopy (TEM) confirmed that the 81R6 strain had an amorphous cell morphology with aberrant division septa, membrane blebbing, and mini-cell formation (Fig 4B). In line with these results, we observed a growth defect for 81R6 compared to *E. faecium* Com12, that could be restored upon complementation (Fig 4C). A similar phenotype was observed for a *sagA* deletion strain of *E. faecium* Com15 (33), suggesting a role of functional SagA for proper cell division. Additionally, 81R6 cells have increased negative cell surface charge which may be a consequence of the elevated cell surface area of clustered unseparated cells (Fig 4D).

**Fig 4.**
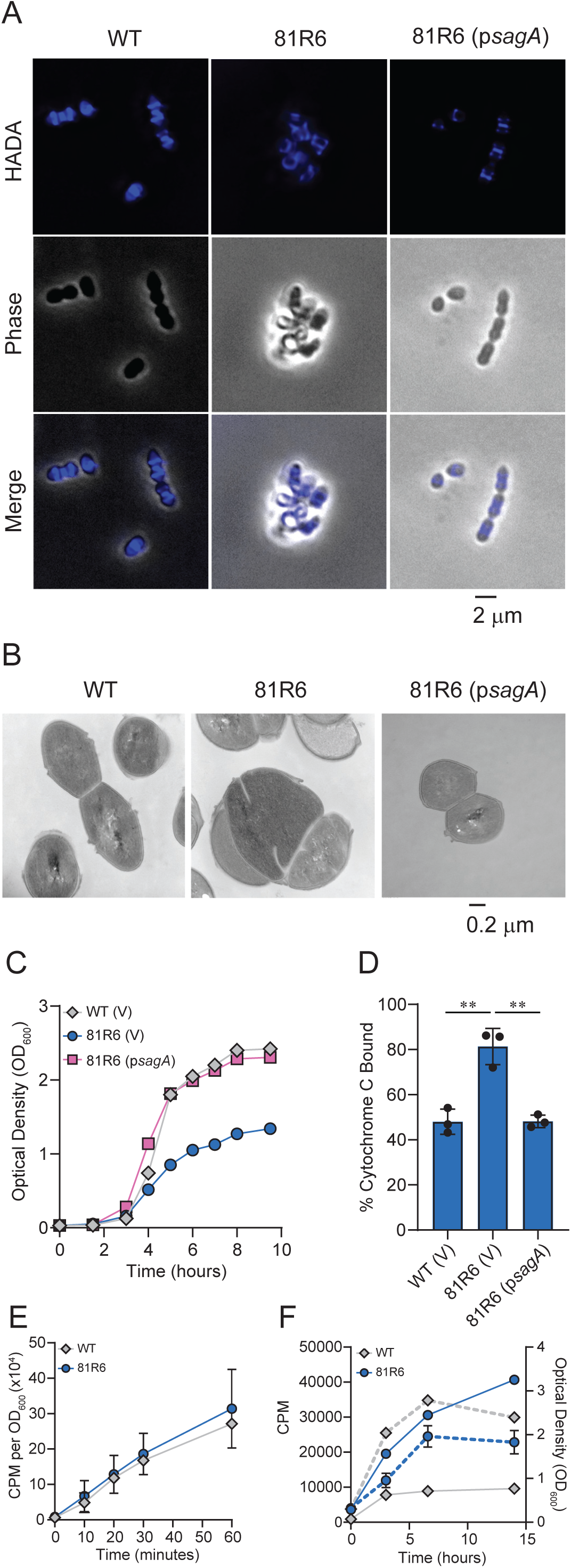
*E. faecium sagA* mutants show aberrant cell division and altered the cell surface charge. **(A)** HADA staining of *E. faecium* Com12 (WT), 81R6, and 81R6 (*psagA*). **(B)** Transmission electron microscopy of *E. faecium* strains shown in panel A. **(C)** Growth curves of *E. faecium* Com12 (WT), 81R6, and 81R6 (*psagA*). V indicates strains carrying the empty vector pAM401. Experiment was performed three times and a representative biological replicate is shown. **(D)** Cell surface charge of *E. faecium* Com12 (WT), 81R6, and 81R6 (*psagA*) using a cytochrome c binding assay. Unbound cytochrome c was quantified in the culture supernatant by measuring absorbance at 570 nm. Values represent mean ± SD of three biological replicates. p values were calculated using unpaired two-tailed Student’s t test. (***p < 0.001; **p < 0.01; *p < 0.03) **(E)** Incorporation of [^14^C]GlcNAc by *E. faecium* Com12 (WT) and 81R6. Values represent mean ± SD of three biological replicates. **(F)** Release of radiolabeled peptidoglycan fragments into the culture medium by *E. faecium* Com12 (WT) and 81R6. Values represent mean ± SD of three biological replicates. Solid lines indicate ^14^C signal measures at counts per minute (CPM) using scintillation counting. Dashed lines indicate growth as measured by optical density (OD_600_).

We next assessed PG synthesis and turnover in *E. faecium* Com12 and 81R6. For PG synthesis we monitored the incorporation of [^14^C] N-acetylglucosamine (GlcNAc) into exponentially growing cells. [^14^C]GlcNAc incorporation increases at the same rate for both *E. faecium* Com12 and 81R6 throughout growth (Fig 4E), indicating that mutation of *sagA* mutation does not affect the rate of PG synthesis under the conditions tested. To evaluate PG turnover, we measured the release of radiolabeled PG fragments into culture supernatants. Cells were grown in media containing [^14^C]GlcNAc, washed, and resuspended in media containing unlabeled GlcNAc. The amount of free radioactivity in the culture media was measured over time. During stationary phase, the release of radioactivity was slow and at a constant rate for *E. faecium* Com12 (Fig 4F). However, 81R6 released PG much more rapidly indicating that more frequent cell turnover attributed to cell lysis in the stationary phage occurs, likely driven by compromised cell envelope integrity (Fig 4F).

### SagA directs phages to adsorb to the cell surface at sites associated with cell division

To adsorb to bacterial cells, phages bind to surface molecules such as polysaccharides, carbohydrate components, and integral membrane proteins that are positioned in spatially distinct locations of the cell envelope (43, 44). Previously, we showed that phages require SagA for infection and that phage adsorption to the *E. faecium* cell surface was unaffected by *sagA* mutation (26). Therefore, it is likely that compromised cell wall integrity and improper cell division in *sagA* mutants alters the spatial distribution of phage receptors that supports non-specific phage adsorption and prevents DNA entry into cells. Thus, we fluorescently labeled phage 9181 with SYBR gold and assessed its distribution at the surface of *E. faecium* and 81R6 cells using fluorescence microscopy. *E. faecium* showed punctate SYBR gold signal at locations that correlated with sites of HADA incorporation (Fig 5A). This indicates that 9181 phages adsorb at or near the sites of active cell division. Interestingly, the majority of 81R6 cells showed faint SYBR gold signal at the cell surface and very intense localized signal at aberrant sites of cell division and near mini cells and membrane blebs (Fig. 5A). We posit that this intense concentrated signal could be due to mis-localized and sequestered phage receptors or phage DNA ejected into mini cell compartments that fail to separate during cell division (Fig. 4B). These differences in phage 9181 phage adsorption were confirmed by transmission electron microscopy showing that *E. faecium* Com12 supported phage adsorption at or near the cell equator and septa (Fig 5B). In contrast, 81R6 cells showed extreme phage aggregation at discreate locations and/or more uniform phage distribution over the surface of the cell (Fig 5B). Equatorial and septal localization of phages was restored by *sagA* complementation (Fig. 5B). Collectively, these results suggest that phage receptors coordinate phage binding and infection near active sites of cell division and are mis-localized upon loss of SagA function.

**Fig 5.**
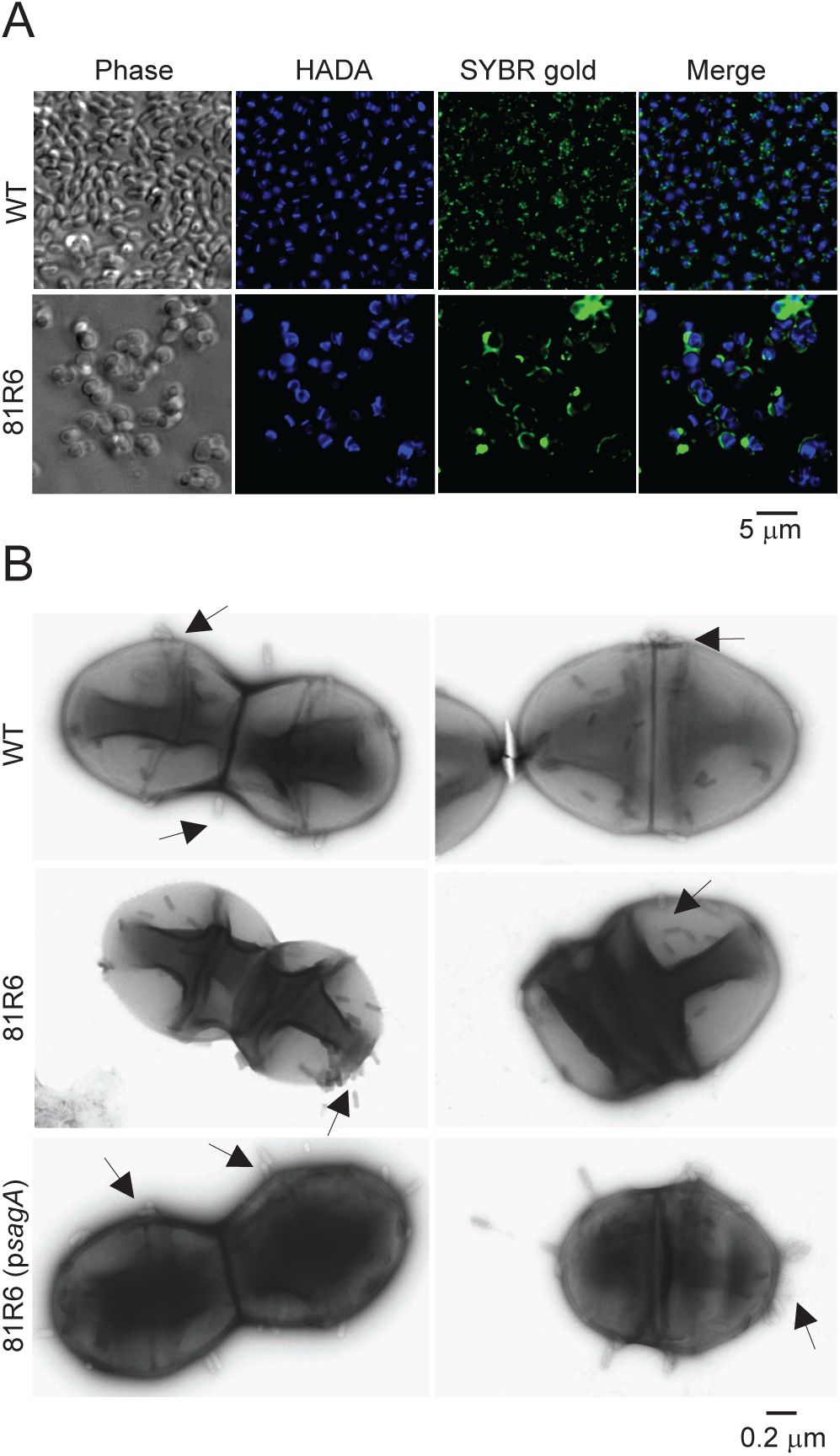
Binding of phage 9181 particles to the surface of *E. faecium*. **(A)** Fluorescence microscopy of *E. faecium* Com12 (WT) and 81R6 shows differential phage adsorption. HADA stained Com12 (WT) and 81R6 cells were mixed with SYBR gold labeled phage 9181 particles. Cells were visualized using fluorescence microscopy 5 minutes post infection. The experiment was repeated two times. Representative images are shown. **(B)** Visualization of phage adsorption on *E. faecium* strains by transmission electron microscopy. *E. faecium* Com12 (WT), 81R6, and 81R6 (p*sagA*) strains were mixed with phage 9181 particles and incubated for 5 minutes at room temperature to allow the phages to attach the bacterial cells. Phage particles are shown with arrows. At least 15 individual images were captured and the experiment was repeated two times. Representative images from single experiment are shown.

## Discussion

Multidrug resistant enterococci and the treatment challenges associated with their infections have renewed attention toward the development of phage therapies. The emergence of phage resistance represents a potential hurdle for successful phage therapies. However, bacteriophage resistance can result in fitness costs, including antibiotic sensitization, colonization defects, and reduced virulence (20, 25, 45–50). Importantly, these fitness costs could be exploited for therapeutic advantage in the clinic, thus extensively characterizing the mechanistic basis of such fitness trade-offs warrants further study. In the present work, we explore β-lactam sensitivity in *E. faecium* lacking functional SagA, a peptidoglycan hydrolase that supports phage infection (26). SagA mutations that support phage resistance lie in the NlpC_P60 hydrolase domain and impair the PG hydrolytic activity of SagA, similar to the previously shown SagA catalytic domain mutants rendering the protein non-functional (31). Non-functional *sagA* results in compromised cell wall integrity, increased cellular permeability, and diffuse distribution of PBPs, rendering cells sensitive to β-lactam antibiotics including cephalosporins. Additionally, *E. faecium sagA* mutants showed irregular distribution of newly synthesized PG in their cell wall and amorphous cell shape. These cell wall synthesis deficiencies further support the observation of cephalosporin sensitivity and show that by altering a critical PG hydrolase, *E. faecium* loses intrinsic cephalosporin resistance. Non-functional SagA also appears to affect the spatial distribution of phage receptors suggesting the role of SagA in mediating phage resistance. To date, detailed studies exploring SagA and its connection with phage and antibiotic resistance in *E. faecium* are limited. Our findings provide insights into the connection between an antibiotic resistance fitness cost resulting from phage selective pressure that could possibly inform approaches to revive cephalosporins as anti-*E. faecium* drugs.

We discovered that *sagA* mutants had cell wall integrity defects and heightened sensitivity to β-lactam antibiotics, specifically cephalosporins. Our data shows that the *sagA* mutant strain 81R6 has a compromised cell envelope with increased permeability to small molecules including β-lactams. β-lactams inactivate PBPs and thereby kill bacteria by inhibiting cell wall synthesis (51). The efficacy of a given β-lactam in inhibiting PG crosslinking usually depends on its reactivity towards the entire repertoire of PBPs expressed by bacterial cells. The increased cell envelope permeability in *sagA* mutants could support rapid PBP occupancy by β-lactams leading to increased killing of *sagA* mutants. We showed that PBP occupancy by Bocillin did not differ significantly between wild type *E. faecium* and the *sagA* mutant 81R6 in the presence of the competing β-lactam ceftriaxone. We did observe, however, that the *sagA* mutant has altered PBP distribution that may affect proper PBP localization and function leading to cell wall synthesis deficiencies. Notably, the high molecular weight PBPs PbpA and Pbp5 are required for intrinsic ceftriaxone resistance in *E. faecium and E. faecalis* (39, 40). A variety of phenotypes similar to those of the *sagA* mutant strain 81R6 are shared with these PBP mutants including impaired cell wall integrity, reduced growth rate, and aberrant cellular morphology (40).

Hydrolases like SagA fulfill cellular PG remodeling demands and maintain the structural integrity of cells during growth. Their disruption can lead to uncontrolled PG degradation and autolysis (52). Tri-septal HADA incorporation into actively dividing *E. faecium* cells suggested a role for SagA in PG remodeling (31). In line with this, 81R6 cells displayed aberrant HADA staining and distorted shape indicating that functional SagA is crucial for proper cell shape and PG remodeling. Specifically, 81R6 showed misplaced septa and bulged cellular morphology indicating improper cell division. PG hydrolases are also required for controlled cell wall synthesis and release PG fragments during cell growth and during PG turnover (52). The *sagA* mutant strain 81R6 showed increased release of radioactive GlcNAC, indicating altered PG turnover. Despite alteration of PG turnover, 81R6 does not have a deficiency in PG synthesis. The increased PG turnover by 81R6 may be attributed to cell lysis due to compromised cell wall integrity and/or increased levels of muropeptides as observed in a similar *sagA* deletion strain (33). Thus, functional *sagA* is required for the controlled release of PG fragments during normal PG turnover.

Bacterial cell growth and division are intimately linked to PG remodeling. PBPs coordinate with the divisome, a complex of proteins involved in PG synthesis, membrane constriction, and cell division. (53–56). In several bacterial species, SEDS-family (shape, elongation, division, sporulation) proteins work with specific PBPs to mediate PG synthesis at specific positions in the cell (57–59). A recent PG synthesis model for enterococci showed that specific PBPs or PG synthases are localized at the site of cell division and elongation that are involved in septal PG and peripheral PG synthesis during growth, respectively (40, 60). Enterococcal PbpA(2b) and Pbp5, which are required for growth in the presence of cephalosporins, are proposed to catalyze PG synthesis at distinct cellular sites (40). Consistent with these findings, our Bocillin labeling experiments showed that PBPs are localized at the equator and septum in wild type cells suggesting their participation in septal and peripheral PG synthesis (Fig. 3C). In contrast, dispersed labeling of Bocillin in the *sagA* mutant 81R6 suggests PBPs are mislocalized leading to growth and division defects. PBPs and cell division proteins work in a coordinated manner for proper PG synthesis. Key cell division proteins interact with different components of the PG synthesis machinery. For instance, FtsZ coordinates with PBPs during Z-ring formation (56) and the FtsEX complex coordinates with PG hydrolases like SagA for proper cell division (61–63). The aberrant septa and unseparated daughter cells observed for 81R6 suggest that the loss of SagA function likely leads to divisome assembly problems. Based on our findings we can infer that SagA works in close association with cell wall synthases, cell division, and elongation machinery and disruption of these interactions likely contributes to the β-lactams sensitivity and phage resistance of *sagA* mutants. However, additional studies are required to further prove a role for SagA in maintaining the fidelity of the divisome and elongasome machinery.

Our study highlights the cost of phage resistance in *E. faecium* resulting in pleiotropic changes in phenotypic traits that promote antibiotic sensitivity. However, this largely depends on the type of phage and how the host responds to the phage imposed selective pressure. In Gram-positive bacteria, the ejection of phage DNA requires penetration of the PG barrier and interaction with the membrane. This is a two-step process where phages first bind to carbohydrate rich molecules to position themselves in proximity of a phage receptor at the cell surface, followed by interaction with the cell membrane (64–66). In enterococci, evidence of this process exists for *E. faecalis* where a transmembrane phage infection protein (PIP_EF_) is required for phage DNA entry but not for the initial phage adsorption that is driven by a surface exposed exopolysaccharide (45, 46). We show here that the phage resistance phenotype in *sagA* mutants is not caused by an adsorption defect but most likely by altered phage receptor localization or modifications to the cell membrane that do not support phage DNA ejection. As *sagA* mutants have cell membrane abnormalities, a membrane embedded receptor could be mislocalized or sequestered to membrane blebs or mini cells that do not support phage DNA entry or replication.

In conclusion, this study provides insights into how phage mediated pressure, which results in phage resistance mutations, alters the cell envelope of *E. faecium* leading to antibiotic sensitization. Our findings reveal that the cell wall hydrolase SagA is critical for successful phage infection and provides insight into the role of SagA in coordinating PBP function that supports cephalosporin resistance. Therefore, phages could be used to select for *E. faecium sagA* mutants that could be more efficiently targeted with cephalosporin therapies. Such a phage-based strategy would be beneficial to help overcome the intrinsic cephalosporin resistance that plagues antibiotic therapies employed against this important nosocomial pathogen.

## Materials and Methods

### Bacterial strains, bacteriophages and growth conditions

Bacterial strains, bacteriophages, and plasmids used in this study are listed in Table S1. *E. faecium* Com12 was cultivated in Brain Heart Infusion (BHI) broth at 37°C with shaking at 220 rpm. *E. coli* strains were propagated in lysogeny broth (LB) at 37°C at 220 rpm. Chloramphenicol was added to media at 10 µg/ml or 5 µg/ml for maintenance of plasmids in *E. coli* and *E. faecium* respectively.

### Antimicrobial susceptibility assay

Minimal inhibitory concentrations (MICs) for each strain were determined using microbroth dilution assays. Overnight grown cells normalized at an OD_600_ of ∼5×10^5^ CFUs were inoculated into 96-well plates containing two-fold serial dilutions of antibiotics in BHI medium (with chloramphenicol for plasmid maintenance when required). Plates were incubated at 37°C with shaking 220 rpm for 24hrs. OD_600_ was measured at different intervals and the lowest antibiotic concentration that showed no growth was noted as the MIC.

### Cell wall integrity analysis

Cell wall integrity was analyzed by testing the susceptibility of intact *E. faecium* cells to sodium dodecyl sulfate (SDS). Briefly, for testing SDS mediated lysis, overnight grown cells were inoculated into BHI medium and grown to exponential phase (OD_600_ ∼0.2). Cells were harvested and resuspended in 50 µl lysozyme buffer (10 mM Tris pH 8.0, 50 mM NaCl, 20% sucrose). Cell suspensions were split into two equal aliquots. One aliquot was treated with 5 mg/ml lysozyme. All the samples were incubated at 37°C for 5 min. SDS Laemmli buffer (1M Tris-HCl [pH6.8), 2% SDS, 50% glycerol, 25% β-mercaptoethanol, 0.02% bromophenol blue) was added, samples were heated at 95°C for 5 min and run on 12% mini SDS-PAGE gels at 110V. PAGE gels was stained with Pierce Silver Stain (Thermo Scientific) according to the manufacturers protocol and visualized using a SYNGENE G:BOX Chemi XX6 gel doc imaging system.

To assess cell envelope permeability via the β-D galactopyranoside (CPRG) hydrolysis assay, *E. faecium* cultures were grown to stationary phase (∼9hrs) in BHI medium supplemented with 40 µg/ml of CPRG (Sigma) at 37°C. Cells were removed by centrifugation and the supernatant was assessed for (CPRG) hydrolysis by measuring absorbance at 570 nm.

### PBP acylation assays

For Bocillin reactivity assays, overnight grown cells were diluted to an OD_600_ 0.03 in BHI medium and grown for 4.5 hrs at 37°C with shaking at 220 rpm. 1 ml aliquots of cultured cells were harvested by centrifugation, washed, and resuspended in phosphate buffered saline (PBS). Cell suspensions were incubated with varying concentrations (0, 0.5, 1, 2, and 4 µg/ml) of Bocillin (Invitrogen) at 37°C for 30 min. Cells were collected by centrifugation, washed and resuspended in PBS. Cells were diluted four-fold and 200 µl was dispensed in triplicate in 96 well plates. Fluorescence was measured using a microplate reader in top mode with excitation and emission at 488 and 517 nm, respectively. Fluorescence was normalized to OD_600_.

For PBP acylation assays, cultures were grown as described above. Cells were treated with ceftriaxone (Sigma) (0, 0.1, 0.5, 1, 8, 64 µg/ml) for 30 min at 37°C with shaking at 220 rpm. Cells were collected by centrifugation at 9400 × *g* for 10 min, resuspended in PBS, normalized to OD_600_ 8.0 in 5 ml and treated with Bocillin 1 µg/ml for 30 min at 37°C. Cells were washed twice with PBS, resuspended in 3 ml of PBS, and disrupted by bead beating using MP Lysing Matrix B tubes (Fisher Scientific) for 10 min by placing samples on ice for 2 min after every cycle. Supernatant was collected by centrifugation at 21130 × *g* for 10 min at 4°C. The membrane fraction was collected by ultracentrifugation at 100,000 × *g* for 30 min. Protein concentration was analyzed by Bradford assay via Bio-Rad protein assay dye reagent concentrate. Samples with equivalent protein concentration were subjected to 10 % SDS-PAGE on 20 cm gel at 140 V for 40 hrs at 4°C. Gels were scanned for Bocillin labeled protein bands at 488 nm using a BioRad Chemi Doc MP Imaging system.

### Microscopy

For visualization of Bocillin stained *E. faecium*, exponential phase (∼0.4-0.5 OD_600_) cells were incubated with 5 µM Bocillin and 0.5 mM HADA (TOCRIS) in BHI media for 30 min at 37°C. Cells were washed twice with PBS and fixed with 1% formaldehyde for 15 minutes at room temperature (RT). Cells were washed and resuspended in PBS. For imaging cells were mounted on 1% agarose pads and imaged under a Zeiss LSM 780 confocal laser scanning microscope.

For HADA labeling, exponentially grown (0.4-0.5 OD_600_) cells were collected by centrifugation at 9400 × *g* and resuspended in BHI. Cells were incubated with 2.5 mM HADA (TOCRIS) at 37°C for 40 minutes, followed by washing with PBS. Cells were fixed with 3.7% paraformaldehyde. Cells were seeded onto 22×22 mm coverslips. For mounting, 10 µl of ProLong Diamond Antifade Mounting Media (Invitrogen) was placed on slides and coverslips laid on slides. Slides were incubated in the dark overnight and imaged using a Nikon Eclipse Ti-2 inverted light microscope equipped with an Orca Fusion BT cMOS camera (Hammamatsu), an IRIS 15 cMOS camera (Photometrics), and Semrock Brightline filters.

To visualize phage 9181 adsorption to the HADA labeled *E. faecium* cells, phage 9181 was purified as described (26). 100 µl of purified phage particles of 10^10^ PFU/ml were mixed with SYBR Gold nucleic acid stain (Invitrogen) at 10,000:5 (V/V) ratio and incubated at RT for 15 minutes. The mixture was washed with SM-plus buffer (100 mM NaCl, 50 mM Tris-HCl, 8 mM MgSO_4_, 5 mM CaCl_2_ [pH 7.4]) three times using 3K Amicon filter (Merck). Exponentially grown *E. faecium* cells (OD_600_ 0.4) were collected, washed and resuspended in BHI and incubated with 2.5 mM HADA for 40 minutes. After two washes cells were resuspended in PBS and mixed with SYBR-gold labeled phages (1:1), incubated for 5 minutes at RT, washed in SM-plus buffer and transferred to poly-L lysine coated microscope slides. Cells were imaged using a Nikon Eclipse Ti-2 inverted light microscope as described above.

For transmission electron microscopy, samples were prepared as described previously (67). Briefly, cells were grown in BHI medium to an OD_600_ of 0.4 and 1.5 ml cells were collected by centrifugation. Cells were washed with cacodylate buffer (CB) (0.1M, pH 7.4) and fixed at 4°C with 150 µl of fixative buffer containing formaldehyde (2%) and glutaraldehyde (2%) in CB. Samples were submitted to CU-Anschutz Electron Microscopy Core Facility for imaging. For negative staining, exponentially grown cells were collected by centrifugation, resuspended in SM-plus buffer and incubated with purified phage 9181 for 5 minutes at RT. Cells were washed with SM-plus, resuspended in fixative buffer and submitted to CU-Anschutz Electron Microscopy Core Facility for imaging. 10 µl of sample was applied to a glow-discharged 300 mesh formvar and carbon coated grid (Electron Microscopy Sciences) for 5 min and blotted with filter paper. The grid was washed one time by applying 10 µl water on the grid and blotted with filter paper. The grids were stained with 10 µl of 0.5% uranyl acetate solution. After blotting, the grids were allowed to dry. Samples were imaged on a Thermo Fisher Tecnai G2 Biotwin TEM (ThermoFisher) at 120 kV with an AMT low mount NS15B sCMOS camera (AMT Imaging)

### Growth curves

Stationary phase cells were inoculated into BHI medium to an OD_600_ of 0.03 and cultures were incubated at 37°C with shaking at 220 rpm. OD_600_ was measured using a Genesys 30 visible spectrophotometer (Thermo Scientific).

### [^14^C]GlcNAc incorporation

Peptidoglycan synthesis was assessed by the incorporation of [^14^C]GlcNAc (PerkinElmer) in exponentially growing cells as described previously with minor modifications (40). Bacteria were grown in BHI medium to an OD_600_ of 0.2 at 37°C with shaking. Cultures were diluted to an OD_600_ of 0.03 in prewarmed BHI medium supplemented with 0.2 µCi/ml [^14^C]GlcNAc. 400 µl aliquots were collected, washed with 1ml PBS, and cell pellets were suspended in 150 µl water. Cells were transferred to 4ml of Econo safe scintillation fluid (Fisher Scientific) and radioactivity was measured by scintillation counting using a Beckman LS6500 scintillation counter. Optical density was measured from parallelly grown label free cells and data is represented as counts per minute per OD_600_.

### Measurement of ^14^C release

Cells were labeled with [^14^C]GlcNAc as described above, collected by centrifugation, washed with PBS, and resuspended in BHI medium containing label free 1mM GlcNAc (Sigma). Aliquots were withdrawn, cells were pelleted, and the amount of released radioactivity in supernatant was measured by scintillation counting.

### Cytochrome c binding assay

Bacterial cell surface charge was analyzed using a previously described method (67). Briefly BHI grown stationary phase cells were collected, washed twice with 20 mM morpholinepropanesulfonic acid - MOPS buffer, pH 7.0, and normalized to an OD_600_ of 7.0. 900 µl of cell suspension was mixed with 100 µl of 5 mg/ml cytochrome c (Sigma) and incubated for 30 min. Cells were pelleted and the absorbance of the supernatant was measured at 530 nm.

## Acknowledgements

This work was supported by the National Institutes of Health grants R01AI141479 (B.A.D.) and (H.C.H.) and American Heart Association Postdoctoral Fellowship 24POST1183502 (G. A.).

